# Biofilm-Derived Curli and Z-DNA Shape Anti-DNA Antibody Responses During Salmonella Infections

**DOI:** 10.64898/2026.03.02.708989

**Authors:** Molly Elkins, Kaitlyn Grando, Courtney Covolo, Diane Spencer, Jacob F. Maziarz, Erin M. Vasicek, Carly DeAntoneo, Francisco J. Albicoro, Shingo Bessho, Zachary W. Reichenbach, Sophia Olubajo, Bettina Buttaro, Siddharth Balachandran, John S. Gunn, David Pisetsky, Çagla Tükel

## Abstract

Antibodies to Z-DNA, a non-canonical DNA conformation with a left-handed zigzag backbone, are abundant in the serum of patients with systemic lupus erythematosus (SLE), with levels increasing with disease activity and flares. As SLE is associated with bacterial infections, and as extracellular DNA (eDNA) within biofilms of several bacterial species has been shown to adopt the Z-DNA conformation, bacterial Z-DNA may represent a source of immunogenic Z-DNA in SLE and other related autoimmune conditions. In these studies, we investigated whether eDNA in Salmonella biofilms also contained Z-DNA and whether such Z-DNA could elicit an antibody response. Using antibody-based staining approaches, we observed abundant eDNA in *Salmonella enterica* serovar Typhimurium (STm) biofilms in both the Z- and canonical B-DNA configurations, consistent with the highly Z-prone nature of the GC-rich *Salmonella* genome. To assess the functional contribution of these DNA conformations to biofilm integrity, biofilms were treated with DNase I, which lacks enzymatic activity against Z-DNA, or with benzonase, a nonspecific nuclease that degrades both B- and Z-DNA. DNase I treatment applied after biofilm maturation was less effective at thinning biofilms than treatment during early biofilm formation, a pattern also observed with benzonase treatment. Purified curli:DNA complexes contained Z-DNA and, when administered intraperitoneally to mice, elicited robust anti-Z-DNA antibody responses. Similarly, infection with invasive STm induced the production of anti-Z-DNA antibodies *in vivo*. Moreover, STm infection in mice fed a diet that promotes biofilm development was associated with increased Z-DNA levels in the cecal lumen and elevated anti-DNA antibody responses. Collectively, these findings suggest that Z-DNA, likely formed by extruded *Salmonella* genomic DNA, and embedded within curli:DNA complexes of STm biofilms, triggers a host immune response and drives anti–Z-DNA antibody production. This work provides mechanistic insight into how bacterial infections and diet-dependent modulation of biofilm formation may contribute to anti-Z-DNA antibody responses in autoimmune diseases like SLE.

## INTRODUCTION

Antibodies directed against DNA are observed in multiple autoimmune diseases; however, in systemic lupus erythematosus (SLE) [1], they are widely used as markers for disease classification and as biomarkers of disease activity [2,3]. In studies on the pathogenesis of SLE, the origin of anti-double-stranded DNA (dsDNA) antibody production has been particularly enigmatic, since immunization of animals with B-DNA, the classic right-handed, Watson-Crick base-paired, double-helical B conformation, fails to induce autoantibody production, even in the presence of carriers and adjuvants [3,4]. In contrast to B-DNA, certain DNA structures, such as Z-DNA, can also induce an immune response when administered to animals [5,6]. Z-DNA is a left-handed helix with a phosphodiester backbone in a zig-zag orientation. The Z-DNA conformation is energetically unfavorable, but alternating purine-pyrimidine sequences can adopt this conformation under certain environmental conditions. The transition from B-DNA to Z-DNA can be facilitated by base methylation or chemical modification (e.g., bromination of bases) [5,7–11]. Z-DNA can potently induce antibodies in animals [11–15], suggesting that, unlike B-DNA, Z-DNA is immunogenic and fails to induce immune tolerance [16]. Although antibodies to Z-DNA are found in SLE patients, the source of immunogenic Z-DNA has remained a mystery for decades, as it had been generally accepted that this type of DNA does not readily occur in nature, at least in mammalian organisms.

DNA is a major component of biofilms, which are multicellular communities of bacteria enclosed in an extracellular matrix (ECM). The ECM of biofilms is composed of nucleic acids, proteins, and carbohydrates that physically protect the bacteria from the environment including the immune system. The secreted protein curli accounts for about 85% of the ECM mass in *Salmonella enterica* serovar Typhimurium (STm) biofilms [17]. STm forms biofilms in the intestinal lumen and on gallstones [18–20]. It is believed that STm strategically employs two lifestyles, planktonic and biofilm, to establish a successful infection [18,21,22]. In mouse models, *Salmonella* infection induces a robust type I interferon (IFN) response [23] and leads to the production of anti-DNA autoantibodies [18,23–25]. Curli becomes hyperinflammatory when complexed with eDNA in bacterial biofilms, eliciting anti-DNA antibodies of the pathogenic IgG2b subtype and generation of type I IFNs and other pro-inflammatory cytokines [18,23,26–29].

Recent studies have demonstrated that bile acids and cholesterol play a central role in promoting *Salmonella* biofilm formation and curli production [19,30,31]. Consistent with these results, a study by Cruz-cruz *et al.* showed that mice fed a high-cholesterol lithogenic diet (LD), which increases intestinal bile acid concentration, exhibited increased STm biofilm formation and curli production in the gut [20]. Notably, Buzzo *et al.* recently showed that eDNA within the ECM of biofilms from *Haemophilus influenzae*, *Pseudomonas aeruginosa*, and uropathogenic *Escherichia coli* adopts a left-handed Z-DNA conformation as a result of interactions with DNA-binding proteins in the ECM [32]. However, whether STm biofilms similarly harbor Z-DNA is unknown; it is also unclear whether biofilm-associated Z-DNA could serve as a source of anti-Z-DNA immune responses during infection.

In this study, we demonstrate that STm biofilms contain Z-DNA, and characterize Z-DNA/curli interactions within the biofilm matrix. We then directly tested whether biofilms or biofilm components drive anti-Z-DNA production by determining whether *in vivo* administration of purified curli complexes containing Z-DNA or oral infection with STm induces anti–Z-DNA antibody production in mice. In parallel, we investigated whether enhancing STm biofilm formation through a lithogenic diet can amplify anti-Z-DNA antibody responses during infection. Together, these findings identify STm biofilms as a key source of Z-DNA, with curli serving as a structural scaffold that facilitates its presentation. Diet-induced enhancement of biofilm formation amplifies these responses, and complementary experiments reveal a mechanistic link between Z-DNA-producing biofilms, bacterial infection, and the development or exacerbation of anti-Z-DNA immune responses.

## RESULTS

### Z-DNA is present in STm biofilms

While Z-DNA can provide structural integrity to certain human pathogenic bacterial biofilms [32], there has been no direct evidence of its role in STm biofilms prior to these experiments. To investigate the presence of eDNA in the Z-conformation in the STm ECM, we grew STm biofilms *in vitro* for 72 hours on glass slides. We performed immunofluorescence staining and imaged biofilms using confocal microscopy. Both B-DNA and Z-DNA were readily detected within the 3-day biofilm ECM, co-localizing with curli in discrete regions enriched for each DNA conformation (Figure 1A). We next examined 7-day-old mature biofilms, as prior studies have shown that Z-DNA content increases as the biofilm matures [32]. We again observed regions enriched for both B-DNA and Z-DNA (Figure 1B). Consistent with previous studies, the biofilms appeared visually thicker at 7 days [32]; however, the relative proportions of B-DNA and Z-DNA did not differ between the 3-day and 7-day time points (Figure 1C). We also noted regions of the biofilm in which Z-DNA and B-DNA were not both present; instead, Z-DNA was observed in the absence of B-DNA (Figure 1D) and in open spaces between cell clusters (Figure 1A). Together, these findings demonstrate that Z-DNA, a non-canonical eDNA, is present throughout STm biofilms and may associate with other ECM components during biofilm development.

**Figure 1.**
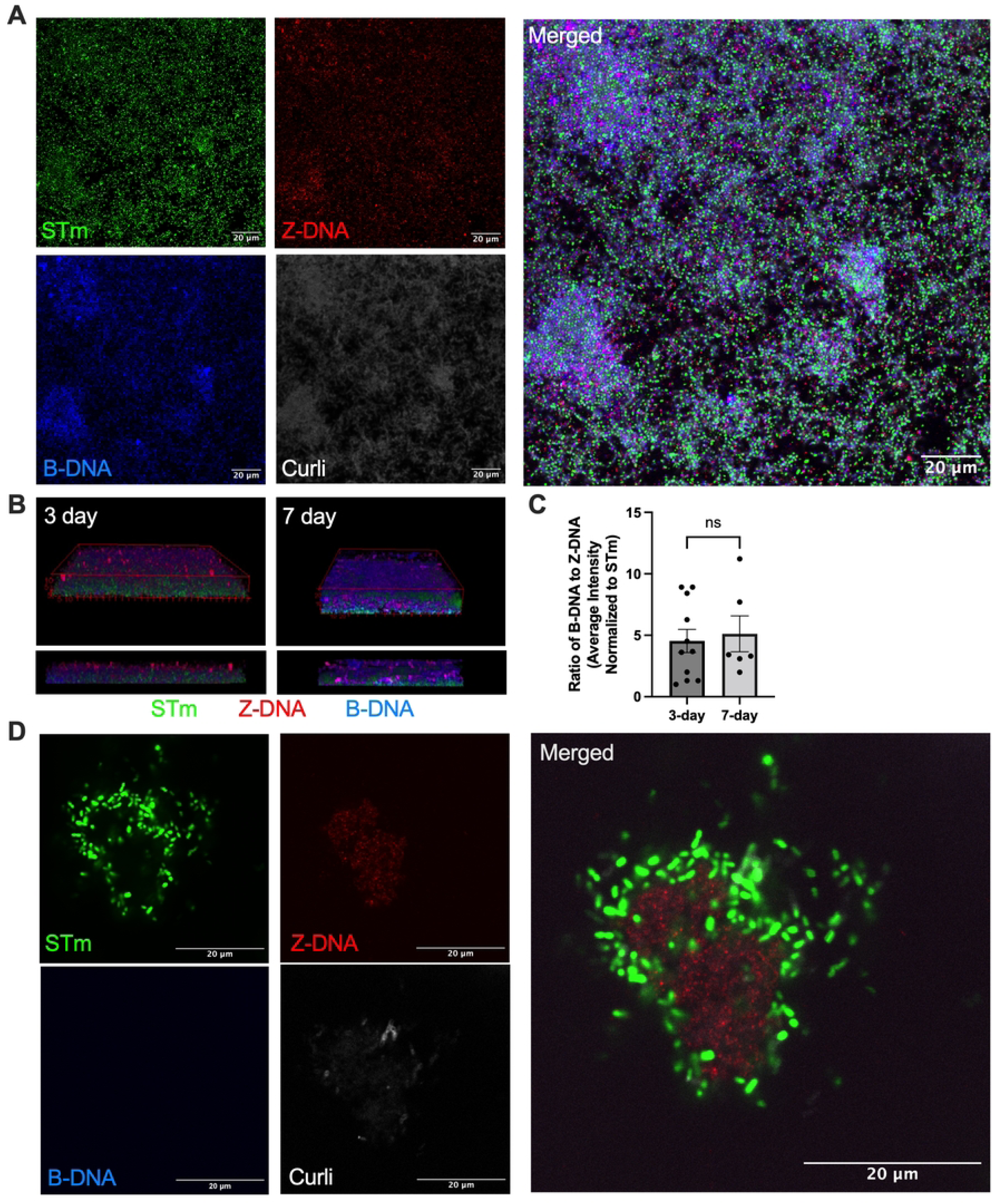
STm biofilms contain Z-DNA. STm was grown on glass cover slips at 28°C for (A-D) 72 hours or (B-C) 7 days and stained with Syto9 dye showing bacterial cells (green), antibodies against Z-DNA (red) and B-DNA (blue), and FSB amyloid dye for curli (gray). Images were acquired at 100X using confocal microscopy and analyzed in ImageJ. Scale bars represent 20µm. (C) The mean fluorescence intensity (MFI) of B-DNA and Z-DNA was measured in ImageJ, the ratio of B- to Z-DNA was calculated, and normalized to the MFI of STm. The experiment was repeated at least three times. Representative data is shown.

### Z-DNA supports structural integrity and modulates biofilm robustness

To further evaluate the contribution of Z-DNA to biofilm structure, we performed DNase treatment assays. Because Z-DNA is inherently resistant to DNase I digestion while B-DNA is readily degraded[33], DNase treatment provides a useful approach to distinguish the proportions of these DNA conformations within the extracellular matrix, and to assess their contributions to the development of the biofilm. To assess the role of different DNA conformations, we added DNase I to STm biofilms at various time points and quantified total biomass using confocal microscopy and Comstat Biomass, which measures 3D physical volume (µm^3^) normalized to area (µm^2^) and biofilm thickness (µm). When DNase I was added at the very start of biofilm development (0 hour), we observed a marked reduction in both biomass and thickness, suggesting that DNase-sensitive eDNA (likely in the B-form) is important for initial biofilm establishment. By contrast, DNase treatment at later stages (24 and 48 hours post-inoculation) produced progressively smaller reductions in these measures (Figure 2A-C). This pattern suggests that as biofilms mature, an increasing proportion of the eDNA adopts the DNase-resistant Z conformation or becomes protected by interactions with other matrix components, such as curli [26].

**Figure 2.**
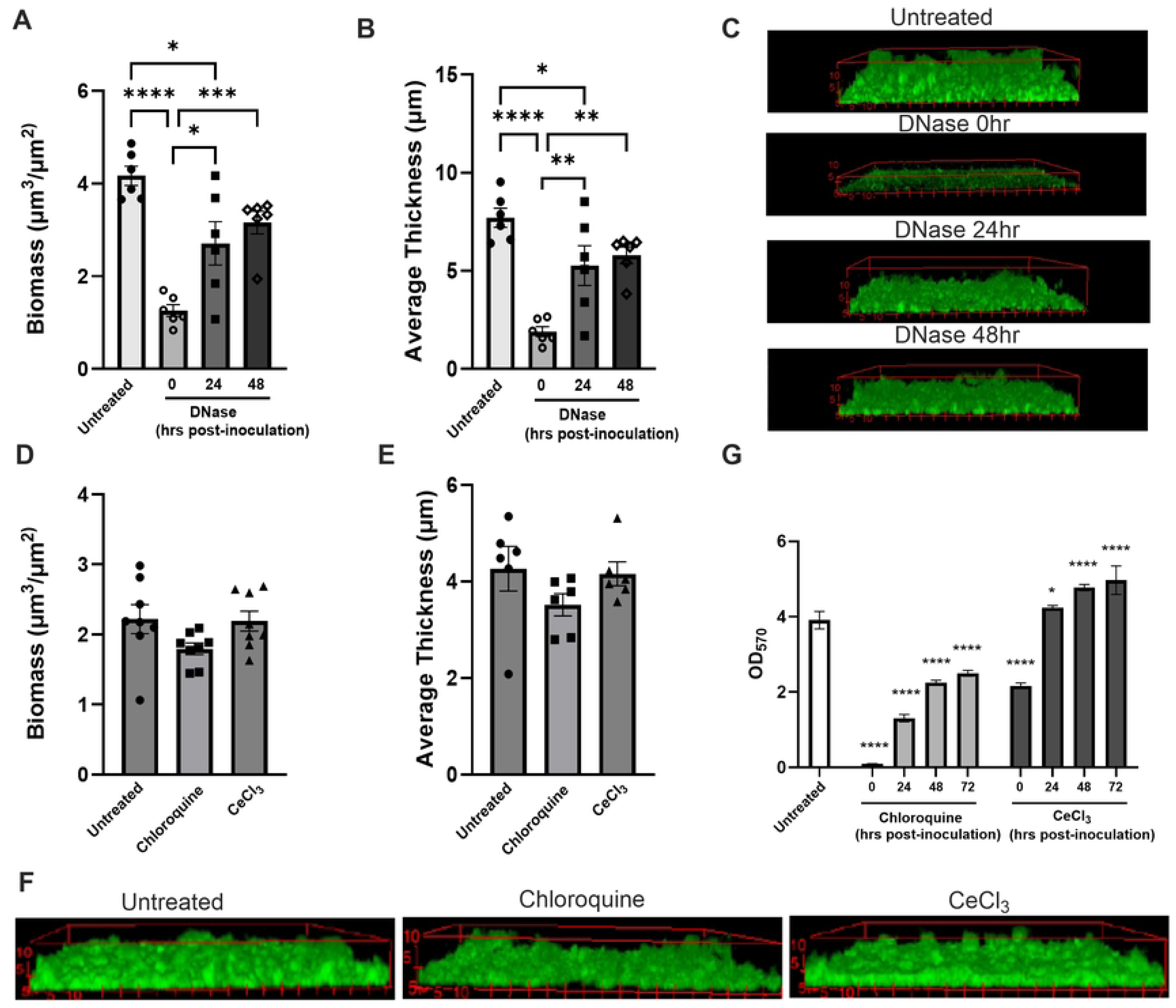
Z-DNA contributes to STm biofilm ECM architecture. (A-C) Biofilms were grown on glass chamber slides and treated with 10U/mL DNase at setup (0 hours), 24 hours, or 48 hours post-inoculation, stained with Syto60, imaged by confocal microscopy at 63X, and analyzed for (A) biomass measured as 3D volume μm^3^ over area μm^2^ (B) average thickness in μm for the entire area via Comstat2 within ImageJ. Confocal images are shown in (C). (D-F) Separate biofilms were treated with chloroquine or CeCl_3_ at 24 hours post-inoculation, then stained with Syto60, imaged via confocal microscopy at 96 hours, and measured for (D) biofilm biomass and (E) average thickness; (F) representative confocal images are shown. (G) Separate biofilms were treated with chloroquine or CeCl_3_ at setup (0 hours), 24 hours, 48 hours, or 72 hours post-inoculation and biofilm robustness was analyzed via crystal violet assay. * represent significance compared to the untreated condition. The experiment was repeated at least three times. Representative data is shown.

Next, we used a previously established chemical approach to shift eDNA toward specific conformations, applying chloroquine to favor the B-DNA conformation and Cerium (III) Chloride (CeCl₃) to promote the Z-DNA conformation [34,35] (Figure 2D-F). We initiated biofilm formation and added chloroquine or CeCl₃ 24 hours after biofilm establishment, then examined biofilm architecture at 96 hours using confocal microscopy and Comstat quantitation. Both treatments showed visual changes in the biofilm architecture. Untreated biofilms were visually homogenous in height with variable cellular packing (density). Chloroquine-treated (B-DNA-enriched) biofilms were visually more variable in height, and exhibited a modest reduction biomass and thickness compared to untreated controls (Figure 2D-F). In contrast, CeCl₃-treated biofilms (Z-DNA-enriched) exhibited a more heterogeneous appearance, with denser bases and structures protruding above the base. However, this altered architecture was not associated with significant changes in biomass or thickness relative to untreated biofilms (Figure 2F). Next, we used the crystal violet assay, which quantifies the amount of stain absorbed to measure cellular density, extracellular components, and charged surfaces, to assess how chloroquine and CeCl₃ influence biofilm biomass when added at different stages of biofilm development. Chloroquine (B-DNA enrichment) was added at 0 h (inoculation) or at 24, 48, or 72 h after biofilm initiation and was maintained in the culture thereafter. In all cases, chloroquine treatment significantly inhibited biofilm formation compared to untreated controls (Figure 2G). In contrast, continuous exposure to CeCl₃ (Z-DNA enrichment) resulted in increased biofilm biomass relative to control conditions (Figure 2G). Notably, even CeCl₃ treatment initiated at 48 or 72 h was sufficient to increase biofilm biomass relative to controls (Figure 2G). Three observations were used for biofilms: visual observations, Comstat-quantified biomass that takes into account the thickness area but does not measure cellular density, and crystal-violet quantified biomass that takes into account the density of cells and ECM. Together, these methods suggest that B-DNA enrichment by chloroquine reduces biofilm density, whereas Z-DNA enrichment by CeCl₃ increases it (visual and crystal violet biomass). Neither treatment significantly changed the 3D space occupied by the biofilm as measured by Comstat biomass and thickness.

Consistent with this model, we next examined how the degradation of extracellular nucleic acids affects STm biofilm structure using benzonase, a broad-spectrum endonuclease from *Serratia marcescens* that cleaves both DNA forms regardless of sequence or conformation [36,37]. Benzonase was applied to developing biofilms at increasing concentrations either at the onset of biofilm formation (0 hours) or after mature biofilms had formed (48 hours post-inoculation). When benzonase was added at the beginning of biofilm development, we observed a reduction in biofilm thickness (Figure 3A–B), indicating that early biofilm assembly depends on intact eDNA. In contrast, the structure of 72-hour established biofilms was less affected by benzonase treatment at 48 hours post-inoculation, suggesting that a portion of the eDNA becomes protected from nuclease digestion as the biofilm matures, potentially through interactions with other ECM components.

**Figure 3.**
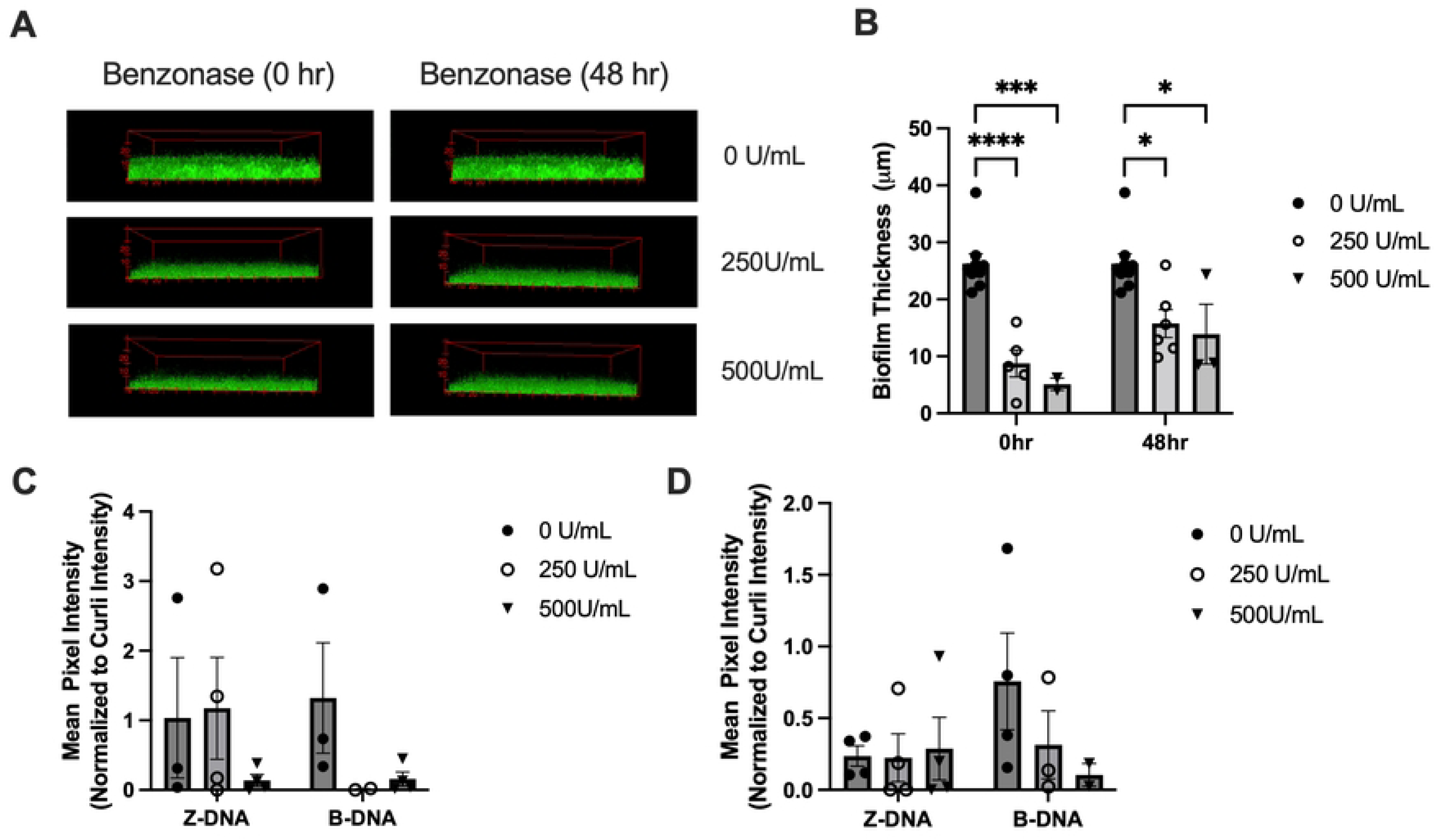
Degradation of Z-DNA and B-DNA reduces biofilm thickness. Biofilms were treated at setup (0 hours) or 48 hours post-inoculation with the indicated doses of benzonase. (A) Biofilms were stained with Syto9, imaged via confocal microscopy at 63X, and (B) biofilm thickness was measured. Biofilms treated with benzonase either (C) at setup (0 hours), or (D) 48 hours post-inoculation were stained for Z-DNA, B-DNA, and curli, and MFI of Z-DNA and B-DNA relative to curli MFI was measured in ImageJ. The experiment was repeated at least three times. Representative data is shown.

To further examine the presence of Z-DNA and B-DNA, we used immunofluorescence to quantify both conformations present in the biofilm matrix following increasing doses (0 U/mL, 250 U/mL, and 500U/mL) of benzonase, at the onset of biofilm growth (0 hours) and after the biofilm was established (48 hours post-inoculation). In both early and established biofilms, benzonase reduced the abundance of extracellular Z-DNA and B-DNA (Figure 3C-D), indicating that both canonical and non-canonical DNA structures are susceptible to enzymatic degradation within the matrix.

Taken together, these findings demonstrate that disruption of biofilm eDNA, including Z-DNA and B-DNA, compromises the structural integrity of STm biofilms. These results further support a model in which Z-DNA contributes to ECM stability, potentially through interactions with other ECM molecules, such as curli, and plays a previously unrecognized role in the maturation and resilience of *Salmonella* biofilms.

### Curli complexes with Z-DNA and induces anti-Z-DNA antibodies

Curli forms amyloid-like fibrils that bind to extracellular bacterial DNA within the ECM of STm biofilms, and these curli:DNA complexes are substantially more immunogenic than either curli or DNA alone [26]. To further define the nature of these complexes, we sought to determine whether curli fibrils preferentially associate with B-DNA, Z-DNA, or both within intact biofilms. To this end, we stained 3-day STm biofilms for B-DNA, Z-DNA, and curli. Consistent with prior observations, curli assembles into basket-like structures that encase individual bacterial cells, appearing as ring-like formations by microscopy [26,38]. As previously reported, the DNA signals were concentrated within these curli rings (Figure 4A). We detected a robust Z-DNA signal throughout STm biofilms, and curli co-localized with both B-DNA and Z-DNA in close proximity to bacterial cells. Notably, Z-DNA appeared tightly associated with curli fibrils within these basket-like structures, appearing as rings around the cells visible in the Z-DNA, whereas B-DNA was more frequently detected at the periphery of curli assemblies, occupying regions that were more exposed to the extracellular environment (Figure 4A). These spatial differences suggest that distinct DNA conformations may differentially interact with curli and contribute uniquely to biofilm architecture and stability.

**Figure 4.**
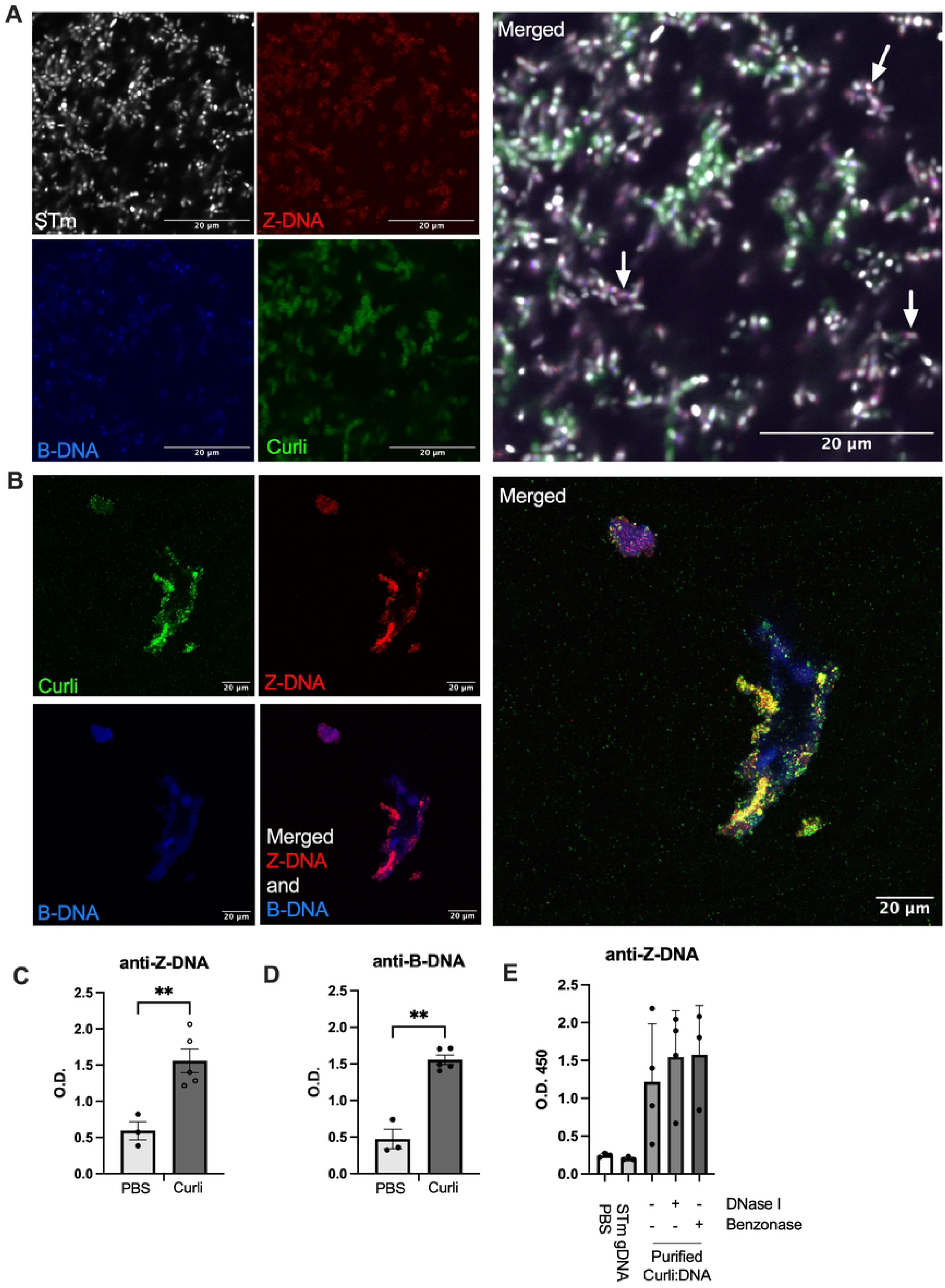
Curli complexes with Z-DNA and elicit anti-Z-DNA antibodies. (A) Biofilms were grown on glass cover slips and stained with Syto9 (gray), anti-Z-DNA antibody (red), anti-B-DNA antibody (blue), and FSB amyloid stain for curli (green). Images were acquired at 100X using confocal microscopy and merged in ImageJ. White arrows indicate examples of colocalization. (B) Purified curli (green) isolated from *in vitro* 72-hour STm biofilms was stained for both Z-DNA (red) and B-DNA (blue). (C-D) C57BL/6 mice were i.p. injected with purified curli:DNA (100 µg), or sterile PBS for controls, weekly for 13 weeks and serum diluted 1:400 was analyzed by ELISA for (C) anti-Z-DNA [brominated poly(dGdC)] and (D) anti-B-DNA (calf thymus DNA) antibodies. (E) C57BL/6 mice were i.p. injected with genomic DNA extracted from STm; purified curli:DNA preparations containing an estimated 50μg curli and 45ng DNA treated with DNAse I, Benzonase, or untreated with nuclease post-purification; or PBS for controls, twice per week for 5 weeks. Serum was diluted 1:400 and analyzed by ELISA for anti-Z-DNA antibodies.

When we purified curli from *in vitro* STm biofilms and used immunofluorescence to visualize DNA, we observed that curli purified from biofilms contains both Z-DNA and B-DNA (Figure 4B). Importantly, the curli purification process includes multiple rounds of DNase I and RNase treatment, boiling in sodium dodecyl sulfate (SDS), and overnight electrophoresis through a preparative SDS-polyacrylamide gel to remove contaminating nucleic acids. The persistence of DNA following these stringent conditions suggests that DNA is tightly bound to and protected by curli.

We have previously shown that intraperitoneal injection of purified curli containing DNA induces the production of antibodies against dsDNA and chromatin within two weeks [26]. Here, we sought to determine whether administration of DNA-containing curli also promotes the generation of antibodies specific to distinct DNA conformations, namely B-DNA and Z-DNA. To address this question, mice were intraperitoneally injected with 100 µg of purified curli weekly for 13 weeks. To distinguish Z-DNA-specific antibodies, we employed the same experimental approach used in prior studies [39,40] to define anti-DNA antibody specificities in mouse sera. Serum antibody responses were assessed by ELISA by coating plates with calf thymus B-DNA and Z-DNA [brominated poly(dGdC)]. Injection of purified curli in mice induced robust antibody responses against both Z-DNA and B-DNA (Figure 4C and 4D).

Notably, in a separate experiment, intraperitoneal injection of STm genomic DNA alone did not result in the generation of anti-Z-DNA autoantibodies, whereas injection of purified curli without any adjuvant, which contains an apparent mixture of Z- and B-DNA, did, indicating that the curli scaffold is required to confer Z-DNA immunogenicity. Furthermore, pretreatment of purified curli with DNase I or benzonase did not reduce the magnitude of the anti-B-DNA or anti-Z-DNA antibody responses (Figure 4D). While DNase I has limited activity against Z-DNA, benzonase degrades DNA regardless of conformation. The lack of any reduction in anti-Z-DNA antibody responses following treatment with either nuclease suggests that DNA associated with curli complexes is physically protected from enzymatic access, limiting effective degradation. Consistent with this observation, DNA extracted from nuclease-treated curli preparations did not show a reduction in DNA concentration compared to untreated samples (data not shown), indicating that these nucleases were ineffective at accessing the DNA within the curli complexes.

### STm genome exhibits a high propensity for Z-DNA formation

While our data demonstrate that Z-DNA is present in STm biofilms and accessible to immune recognition, its origin remains unclear. To address this question, we analyzed the STm genome using the previously established Z-HUNT and Z-DNABERT algorithms, which predict the energetic favorability of DNA sequences to adopt the left-handed Z-DNA conformation [41]. This analysis revealed that the STm genome exhibits a high number of ZH-sites per 1KB (∼733-713), indicating a strong intrinsic tendency to form Z-DNA structures (Figure 5A, B). These results are consistent with the GC-rich nature of the *Salmonella* genome (∼52.3% GC content [42,43] 44.3%-36.7% for human, mouse, rat, or ape mitochondrial genomes [44]). Notably, the genome of the closely related bacterium *E. coli* also displayed 702-691 ZH-sites per 1KB, and a high corresponding GC content (∼50.6% GC content, Fig 5A,B). In contrast, common gut commensals, including *Bacteroides fragilis* and *Lactobacillus rhamnosus*, showed substantially lower Z-DNA-forming potential (Figure 5A, B). It is important to note that these also show 43.1% and 46.6% GC content, respectively. These results together show a strong correlation between GC-rich genomes and the propensity to form Z-DNA (Spearman’s ρ = 0.98), a trend well established [45,46]. Interestingly, *Mediterraneibacter gnavus* (formerly *Ruminococcus gnavus*), a bacterium whose expansion in the gut of SLE patients also positively correlates with disease severity [45] did not exhibit highly Z-prone genome or contain a GC-rich genome (42.8%), suggesting that Z-DNA propensity is not a universal feature of inflammation-associated microbes.

**Figure 5.**
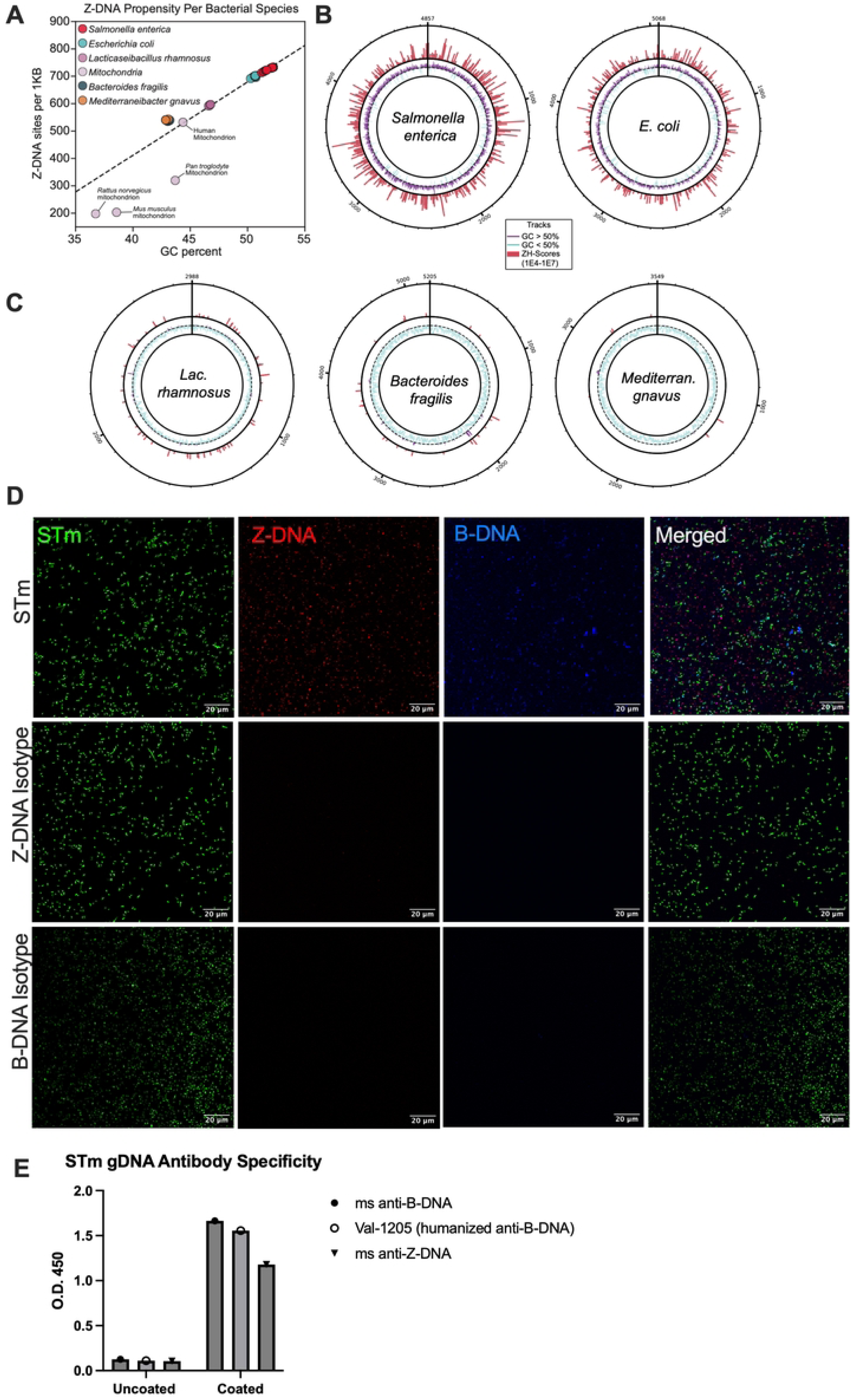
The STm genome contains regions in Z-configuration. (A) Five bacterial genomes and 4 mitochondrial genomes were analyzed using the ZHUNT algorithm, and plotted based on Z-DNA sites per 1KB (Y-axis) and GC% (X-axis). Genome-wide Z-DNA–forming potential visualized using ZDNABERT-filtered ZHUNT scores. . Circos plots show (B) *Salmonella enterica* serovar Typhimurium str. LT2 and *Escherichia coli* str. K-12 substr. MG1655; and (C) *Bacteroides fragilis* NCTC 9343*, Lacticaseibacillus rhamnosus* NCTC 13764, and *Mediterraneibacter gnavus* ATCC 29149. The outer red track indicates ZH-scores ranging from 1×10⁴ to 1×10⁷, representing increasing Z-DNA-forming propensity. The inner track shows GC content, with regions >50% GC shown in purple and <50% GC shown in blue. (D) STm (green) was lysed during log phase growth, fixed, and stained for STm-exposed nucleic acids, including Z-DNA (red) and B-DNA (blue). (D) Plates were coated with STm gDNA, and binding of mouse (ms) anti-Z-DNA, mouse (ms) anti-B-DNA, and humanized anti-B-DNA (Val-1205) antibodies was analyzed by ELISA.

To experimentally validate whether STm genomic DNA (gDNA) contains regions capable of adopting the Z conformation, we lysed STm cells during logarithmic growth, fixed the lysates to maintain supercoiling, and stained the exposed genomic DNA with Syto9 or antibodies specific for Z-DNA and B-DNA. As these experiments indicated, both Z- and B-DNA-specific antibodies robustly bound to the lysed STm genome, indicating the presence of both DNA conformations within bacterial chromosomal DNA (Figure 5D). Consistent with these findings, both anti-Z-DNA and anti-B-DNA antibodies bound efficiently to purified STm gDNA, as tested by coating plates with STm gDNA and analyzing antibody binding via specific ELISA assays (Figure 5E). Together, these findings demonstrate that the STm genome contains intrinsic sequence features that favor Z-DNA formation, supporting the conclusion that bacterial genomic DNA is a plausible source of the Z-DNA detected within curli-associated biofilm complexes.

### *In vivo* STm infection induces the generation of anti-Z-DNA antibodies

To investigate how a STm infection affects the generation of Z-DNA antibodies, we orally infected 129X1/SvJ mice for 13 weeks with wild-type STm. These mice developed robust anti-Z-DNA antibody responses. In contrast, infection with a non-invasive *invAspiB* mutant failed to induce detectable anti-Z-DNA antibodies (Figure 6A). Notably, wild-type STm infection also elicited anti-B-DNA antibodies, but these responses were lower at 13 weeks post-infection than those directed against Z-DNA (Figure 6B). Together, these findings demonstrate that, once bacteria breach the intestinal epithelial barrier, invasive STm infection may drive a strong and preferential anti-Z-DNA antibody response.

**Figure 6.**
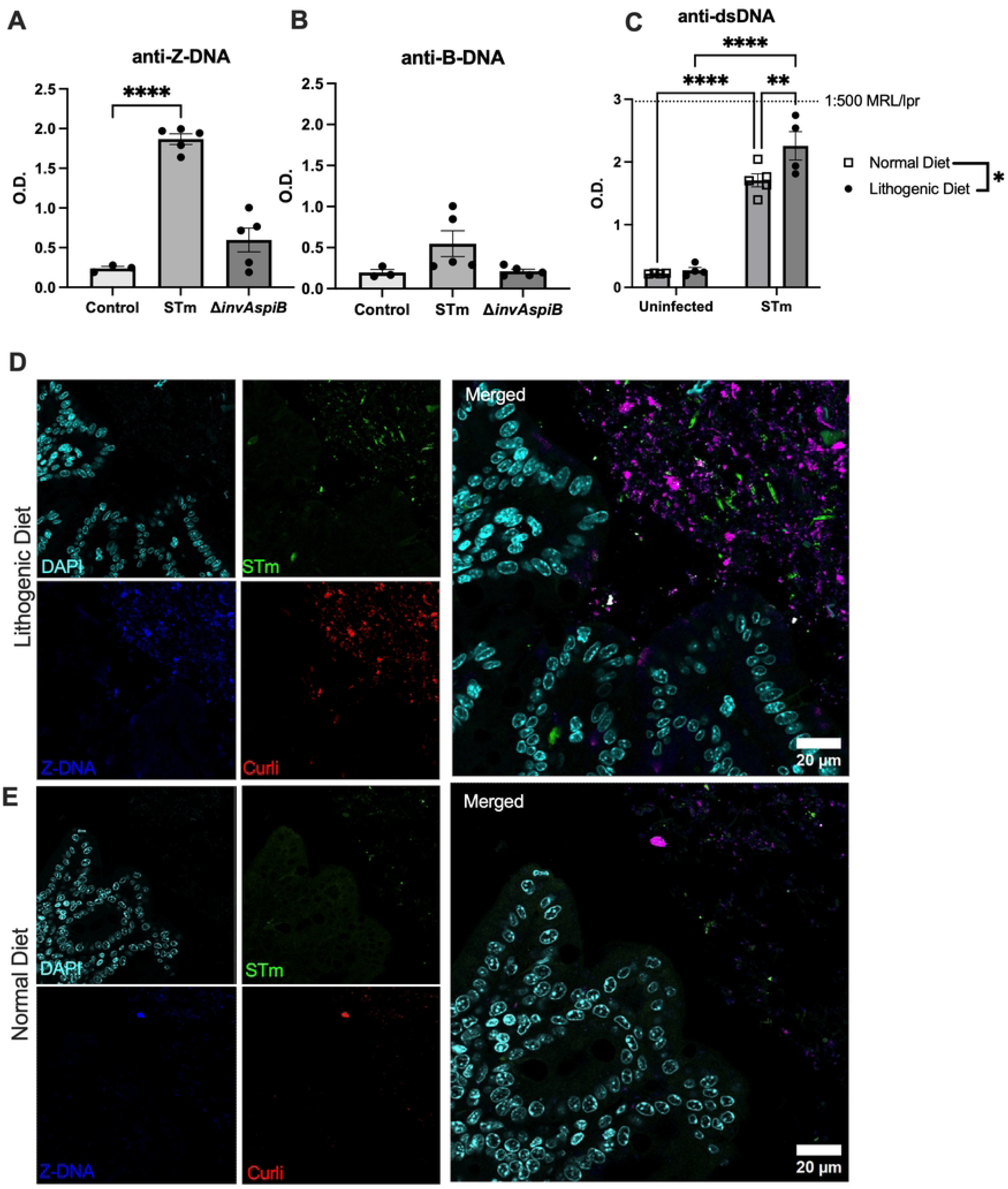
A lithogenic diet induces Z-DNA-containing STm biofilms and higher anti-DNA antibodies. (A-B) 129X1/SvJ mice were intragastrically infected with STm IR715 or an isogenic non-invasive mutant STm (*invAspiB*). The serum was collected 13 weeks post-infection, diluted 1:600, and assayed by ELISA for (A) anti-Z-DNA and (B) anti-B-DNA antibodies. (C-E) 129X1/SvJ mice were fed a lithogenic (high-cholesterol) diet for 8 weeks, then switched to a normal diet for 1 week before i.p. infection with STm. (C) Serum from 3 weeks post-infection was diluted 1:250 and analyzed by ELISA for anti-dsDNA autoantibodies. (D-E) Cecal tissue was stained with antibodies against *Salmonella* (green) and Z-DNA (blue), EBBA Biolight 680 bacterial amyloid dye for curli (red), and DAPI for host nuclei (cyan). Representative images were taken at 63X magnification on the confocal microscope and processed via ImageJ.

Recent work has shown that a high-cholesterol lithogenic diet (LD; 0.5% w/w cholesterol), commonly used to induce hypercholesterolemia in mice, enhances STm biofilm formation in the gut, which correlates with elevated cecal cholesterol level[20]. To determine whether increased biofilm formation influences autoantibody responses, 129X1/SvJ mice were fed LD for 6 weeks and then switched to normal chow (normal diet, ND) prior to i.p. infection with wild-type STm for 3 weeks. LD-fed mice developed significantly stronger anti-dsDNA autoantibody responses than mice fed ND (Figure 6C). Consistent with these serologic findings, histological analysis of intestinal sections from LD-fed mice revealed that STm biofilms exhibited an abundance of Z-DNA in LD-fed mice, and that biofilms appeared closer in proximity to the epithelial surface compared to ND-fed mice (Figure 6D-E). These data support the idea that diet-induced changes in biofilm architecture and localization enhance exposure to immunostimulatory Z-DNA and promote autoantibody production

## DISCUSSION

Infection has long been suspected to contribute to the induction of anti-DNA antibodies, yet the underlying mechanisms remain incompletely defined. The findings presented here demonstrate that bacterial biofilms, specifically biofilm-associated DNA complexed with curli, constitute a potent, structurally distinct source of immunogenic DNA that drives anti-DNA antibody production. These data indicate that both bacterial DNA and bacterial products such as curli can promote anti-DNA responses in normal immunity and may contribute to the pathogenesis of SLE, the human disease most strongly associated with aberrant immune responses to DNA [3,39,40,46]. Anti-DNA antibodies are observed in multiple autoimmune diseases, including SLE, rheumatoid arthritis, Sjögren’s syndrome, and autoimmune hepatitis [47]; however, in SLE they correlate particularly well with disease activity and clinical flares. Indeed, SLE is considered the prototypic autoimmune disease in which anti-DNA antibodies serve as a defining serologic hallmark. These antibodies display features of antigen-driven selection, including somatic hypermutation and affinity maturation, consistent with a sustained and specific immune response to DNA-containing antigens [46,48]. Although host-derived DNA released during apoptosis and necrosis has long been considered a major source of immunostimulatory nucleic acids in SLE, our findings identify bacterial biofilms as an additional, structurally distinct reservoir of immunologically active DNA.

Our findings provide a useful perspective on the roles of both infection and the microbiome in lupus pathogenesis. Several studies have now reported that Pseudomonadota (formerly Proteobacteria), the dominant phylum of Gram-negative bacteria, which includes the *Enterobacteriaceae* family containing *Salmonella* and *E. coli*, as well as the Gram-positive *Mediterraneibacter* (formerly *Ruminococcus*) are overrepresented in the intestinal microbiota of patients with active SLE [51–53]. However, the mechanisms by which these bacteria contribute to pathogenesis remain largely unknown. Using STm as a model organism, we provide evidence for a unique mechanism by which Proteobacteria-derived biofilms may promote SLE-relevant immune responses to DNA, supporting the role of both foreign and self DNA as triggers for autoantibody production.

Using *in vitro* model systems, our studies demonstrate that STm biofilms contain abundant eDNA in both the canonical B-DNA and non-canonical Z-DNA conformations, with levels increasing as the biofilm matures (Figure 1). Under optimal biofilm-forming conditions, curli expression is upregulated over time and reaches maximal levels at approximately 48 hours, coinciding with advanced matrix development. eDNA is a well-established structural component of the biofilm ECM, contributing to biofilm integrity and stability. Importantly, our functional analyses reveal that both B-DNA and Z-DNA are required to establish biofilm architecture (Figures 2 and 3), indicating that the transition of DNA into the Z-conformation actively contributes to biofilm structure rather than representing a passive byproduct of biofilm maturation. Within the ECM, DNA is highly organized and complexed with matrix-associated proteins. Previous work has shown that DNABII family proteins play a critical role in the initial stabilization and organization of eDNA in biofilms; however, DNABII-associated DNA remains sensitive to nuclease degradation [32,54–57]. Although the precise role of curli in relation to biofilm DNA has not been fully defined, our data support a model in which curli interacts with DNA following its initial organization by DNABII proteins, thereby further stabilizing the biofilm matrix. Notably, the β-sheet motif of curli monomers organizes DNA into a spatially periodic lattice that amplifies immune activation [27,58]. In addition, DNA enhances curli polymerization [26], indicating a reciprocal interaction that promotes amyloid scaffold assembly within the ECM. Once curli is polymerized and incorporated into the matrix, DNA associated with curli becomes markedly more resistant to nuclease degradation (Figures 3B–D, 4E), demonstrating that the amyloid scaffold both reinforces and protects the ECM.

Consistent with this model, curli-associated DNA is displayed to the immune system in a highly organized manner, forming an ordered and multivalent array of parallel dsDNA. This structured presentation enhances engagement with innate immune receptors, including TLR9, thereby promoting immune activation [27,58]. Curli:DNA complexes, but not purified genomic STm DNA alone, induced antibody responses *in vivo* (Figure 4C-D). Although these complexes contain both B-DNA and Z-DNA (Figure 4B), i.p. injection elicited both anti-B-DNA and anti-Z-DNA autoantibodies, demonstrating that curli association is essential for DNA immunogenicity. Notably, although *in silico* analysis using the Z-Hunt algorithm predicts that the GC-rich STm genome is highly prone to adopting the Z-DNA conformation, genomic DNA alone failed to induce anti-Z-DNA antibodies (Figures 4E, 5A). These findings underscore the importance of curli-mediated stabilization and presentation of Z-DNA for elicitation of an effective anti-Z-DNA antibody response. In this context, curli likely functions as both a carrier and a structural scaffold, converting DNA from a normally inert molecule into a potent danger signal. We have previously shown that curli:DNA complexes engage both TLR2/TLR1 and TLR9 signaling pathways [23,59–62], further supporting a role for curli as a delivery system that promotes innate immune sensing and downstream autoimmune responses.

Importantly, differences in GC composition across bacterial genera influence TLR9 activation, as this receptor specifically recognizes unmethylated CpG-containing DNA motifs. Indeed, prior analyses of DNA from multiple bacterial species demonstrated variable TLR9-activating capacity, with *E. coli* exhibiting both the highest CpG content and the strongest activation[63]. Beyond CpG frequency alone, genomic propensity to adopt alternative DNA conformations may further modulate immunogenicity. Comparative Z-Hunt analyses revealed that STm and closely related *E. coli* genomes are highly enriched in regions with a propensity to adopt the Z-DNA conformation, whereas *R. gnavus*, a gut commensal associated with SLE, exhibits a lower Z-DNA–forming potential (Figure 5A). This analysis suggests that members of the phylum Proteobacteria may be inherently better able to generate Z-DNA–rich extracellular matrices compared to other commensals and *R. gnavus*, particularly when DNA is present in curli-containing biofilms. As a result, Proteobacteria-derived biofilms may represent a more potent source of immunogenic Z-DNA than those of *R. gnavus*, thereby driving more effective anti-Z-DNA antibody responses in SLE patients. Together, these findings support a model in which the taxonomic composition of the gut microbiota, including the enrichment of Z-DNA–prone Proteobacteria, influences an array of microbial signals that act on the immune system and drive anti-DNA production. Extending these observations *in vivo*, we find that invasive STm infection induces anti–B-DNA antibodies; however, these responses are modest and significantly lower than those directed against Z-DNA. In contrast, non-invasive STm infection confined to the intestinal lumen fails to elicit anti–DNA antibody responses (Figure 6A), indicating that luminal exposure alone is insufficient to break immune tolerance. These data demonstrate that invasive STm infection, when bacteria breach the intestinal epithelial barrier and gain access to host immune cells, drives a robust and preferential anti–Z-DNA response [64,65].

To further elucidate the amplification of these responses by host factors (e.g., diet), we employed a lithogenic (high-cholesterol) diet that promotes bile acid–dependent biofilm formation in the cecum [20]. Use of this diet resulted in a marked increase in anti-DNA antibody production during STm infection (Figure 6C), indicating that diet-enhanced biofilm formation can amplify systemic anti-DNA responses. In LD-fed mice, STm biofilms extended closer to the epithelium and exhibited significantly greater Z-DNA abundance compared to mice fed a standard diet (Figure 6D-E). These findings suggest that dietary modulation of the gut environment can promote biofilm formation and boost the immune responses to biofilm-associated molecular patterns.

Beyond SLE, invasive enteric infections, including STm, are clinically associated with reactive arthritis (ReA) [66–72]. While most individuals experience self-limiting gastroenteritis, a subset, particularly those carrying the HLA-B27 genotype, develop chronic inflammatory arthritis weeks after infection [73–76]. In mouse models, STm biofilms and curli produced in the intestine play critical roles in joint inflammation and synoviocyte hyperplasia [18], further supporting the idea that immune signaling by biofilm molecules can drive chronic inflammation. A relevant question is whether antibody responses to DNA are similarly observed in patients with ReA and whether these antibodies contribute to the disease.

In addition to demonstrating the influence of infection on anti-DNA production, this study shows that dietary changes can alter the organization of the gut microbiota and its biofilm-forming capacity, thereby affecting the quality and quantity of microbial signals that influence the immune system. By promoting Z-DNA-rich biofilms, dietary factors, such as a lithogenic diet, may amplify anti-DNA antibody responses. In the future, it will be critical to determine whether bacterial biofilms and biofilm-derived nucleic acid/protein complexes play a broader role in promoting inflammation in SLE and other conditions, potentially linking intestinal dysbiosis, autoantibody production, and systemic inflammation.

## MATERIALS AND METHODS

### Bacterial growth

*S.* Typhimurium strain 14028 and *S.* Typhimurium strain IR715 were used in this study. *S.* Typhimurium strain IR715 is a fully virulent, nalidixic acid-resistant strain derived from the ATCC strain 14028[77]. The SPI-1/SPI-2 type three secretion-negative IR715 Δ*invAspiB* mutant was described previously [78,79]. Single colonies of STm were inoculated into 5 mL Luria Broth (LB) supplemented with 50 μg/mL nalidixic acid or Tryptic Soy Broth (TSB) and grown overnight at 37°C with shaking.

### In vitro biofilms

Sterile glass coverslips were placed into a 24-well Fisherbrand™ Tissue Culture Plate (FB012929), and the surrounding wells were filled with sterile water to prevent drying. STm cultures grown overnight were diluted 1:100 in LB without salt (LBNS) to 300 μL per well. The plate was covered and incubated statically at a 45° angle in a 28°C incubator for each time point (72 hours or 7 days) without media changes. The supernatant was discarded, and each well was gently washed three times with sterile PBS (sPBS). Biofilms were blocked for 1 hour with a 1:200 dilution of Anti-CD16/CD32 FC Shield Antibody and then washed with sPBS. Biofilms were incubated at room temperature for 1 hour with primary antibodies (5μg/mL), washed, then incubated for 1 hour with 1:200 dilution of secondary antibodies. Slides were incubated at room temperature for 15 minutes with Syto9 Green Fluorescent Nucleic Acid Stain (0.3%), then washed with sPBS. Glass coverslips were carefully removed from wells and placed top-down on an 8-well microscope slide containing a drop of Vectashield. Coverslips were sealed with clear nail polish and imaged at 100X using a Leica SP5 confocal microscope. The following commercial antibodies were used for immunofluorescence: 5μg/mL rabbit IgG isotype control (AC042), 5μg/mL mouse IgG2b isotype control (02-6300), 5μg/mL mouse anti-Z-DNA/anti-Z-RNA antibody [Z22] (Absolute Antibodies Ab00783-3.0) (Z-DNA), 5μg/mL rabbit anti-dsDNA antibody [DSD/4054R] (Novus Biologicals NBP3-07302-100ug) (B-DNA). The following secondary antibodies were used in a 1:200 dilution: AlexaFluor555-conjugated AffiniPure donkey anti-mouse IgG (H+L) (0.50mg) (Jackson Immuno Research 715-565-150) and goat anti-rabbit IgG H&L (Alexa Fluor® 647) (Abcam ab150079).

STm biofilms were grown in LBNS and treated with varying concentrations of benzonase (0U/mL, 250U/mL, 500U/mL) at the beginning of biofilm growth (0 hours) or after maturation (48 hours post-inoculation). Biofilms were stained at room temperature for 15 minutes with Syto9 Green Fluorescent Nucleic Acid Stain and mounted and imaged as described above.

STm biofilms grown in 1:20 TSB media were treated with DNase (10U/mL, ThermoFisher), chloroquine (0.5 mM, Sigma Aldrich), or CeCl_3_ (0.5 mM; gift from the Goodman Lab at Nationwide Children’s Hospital). DNAse, chloroquine, or CeCl_3_ were added with the media and the inoculum (0 hours), or at 24 hours, 48 hours, or 72 hours post-inoculation, before analysis at 96 hours. Media containing the appropriate treatment was replaced daily. After 96 hours, the biofilms were washed once with PBS. Cells were labeled with Syto60 (5 mM; Molecular Probes) in 5% bovine serum albumin (BSA) blocking buffer at room temperature for 30 minutes, after which time the stain was carefully removed and discarded. The wells were washed three times with PBS before the addition of 200 µL 4% paraformaldehyde (PFA; Affymetrix) at room temperature for 20 minutes. Stained biofilms were visualized, and three-dimensional biofilm images were acquired by capturing 2 random Z-stacks per well, 3 wells per treatment, using a Zeiss LSM 800 confocal laser scanning microscope at 63X magnification. The Z-stacks were then analyzed using the software package Comstat2 to calculate biomass as 3D volume (μm^3^) over area (μm^2^) and average thickness (μm) for the entire area.

### *In vitro* biofilms for crystal violet assay

A single colony *S.* Typhimurium 14028 was grown in 5 mL TSB at 37°C overnight on a roller drum. The overnight culture was normalized to OD_600_ = 0.47 (optical density at 600 nm) and diluted 1:2500 in 1:20 TSB. After diluting, 200 µL of the diluted culture was added to a non-treated polystyrene 96-well plate in quadruplicate. The plate was incubated for 96 hours at 25°C statically, with 1:20TSB media containing the appropriate treatments (untreated, 0.5mM chloroquine, 0.5mM CeCl_3_) replaced every 24 hours. At 96 hours, the plate was washed twice with double-distilled water (ddH_2_O) and heat-fixed for 1 hour at 60°C. The plate was stained with 0.33% crystal violet solution for 5 minutes at room temperature. After two washes with ddH_2_O, 100 µL of 33% acetic acid was added. The OD_570_ of was measured in a SpectraMax spectrophotometer with SoftMax Pro software (Molecular Devices) to determine the remaining amount of crystal violet stain. This was performed in triplicate.

### Curli purification

Curli aggregates were purified from the *S.* Typhimurium IR715 *msbB* mutant as previously described [80]. After the purification steps, curli preparations were then resuspended in sterile water. Concentrations of curli aggregates were determined using the bicinchoninic acid (BCA) assay according to the manufacturer’s instructions (Novagen, 71285–3). Curli protein preparations were adjusted to be 1 mg/mL, aliquoted and stored frozen at −20°C.

### Immunofluorescent imaging of purified curli

8-well microscope slides were coated for 1 hour with 50ul poly-L-lysine. The slides were washed with sPBS, and 50 μL of 1 mg/mL curli protein preparation was added to each well, and incubated overnight at 4°C. Wells were washed with sPBS and incubated with the antibodies named below. Glass coverslips were placed on top of each well and were sealed with clear nail polish and imaged using a Leica SP5 confocal microscope.

The following commercial antibodies were used: 5μg/mL rabbit IgG isotype control (AC042), 5μg/mL mouse IgG2b isotype control (02-6300), 5μg/mL mouse anti-Z-DNA/anti-Z-RNA antibody [Z22] (Absolute Antibodies Ab00783-3.0) (Z-DNA), 5μg/mL rabbit anti-dsDNA antibody [DSD/4054R] (Novus Biologicals NBP3-07302-100ug) (B-DNA). The following secondary antibodies were used in a 1:200 dilution: AlexaFluor555-conjugated AffiniPure donkey anti-mouse IgG (H+L) (0.50mg) (Jackson Immuno Research 715-565-150) and goat anti-rabbit IgG H&L (Alexa Fluor® 647) (Abcam ab150079).

### Treatment of mice with purified curli

Male and female C57BL/6 (wild type) mice were purchased from Jackson Labs at 4-6 weeks old. At 6–8 weeks of age, mice were injected intraperitoneally (i.p.) with 100 µg of curli:DNA complex or sterile PBS (control) once per week, alternating sides for 13 weeks. After euthanasia with CO_2_, mice were exsanguinated and the blood was collected for analysis.

For the experiments using nuclease-treated purified curli, aliquots of 50 µg curli:DNA complexes were either untreated (control) or incubated at 37°C for 30 minutes with 500 U/mL DNase I or 500 U/mL Benzonase (Sigma-Aldrich, 9025-65-4). Treatments were then incubated at 80°C for 30 minutes to denature the nucleases. Mice were i.p. injected with 100 µL PBS, 45 ng STm genomic DNA as controls, or 50 µg of curli:DNA preparations twice per week for 5 weeks [27].

### Anti-DNA ELISA

The ELISA to quantify anti-dsDNA antibodies was performed as previously described [39,40]. Z-DNA and B-DNA antigens were obtained and prepared as previously described [39,81]. Briefly, poly(dGdC) was brominated (Br-poly(dGdC)) to serve as the Z-DNA antigen according to the following protocol: poly(dGdC) was reconstituted with TE buffer, the sodium chloride content of an aliquot of this stock was adjusted to 150 mM, and the stock was then diluted to 200 μg/mL with citrate/EDTA/NaCl buffer. Bromine water, diluted 1:25 with UltraPure Distilled Water, and poly(dGdC) were mixed in a 1:3:1 ratio and incubated for 20 minutes in the dark at room temperature. Commercially available calf thymus DNA (Sigma-Aldrich) served as the B-DNA antigen.

ELISA assays were performed as described previously [39]. Briefly, plates were coated overnight with 100 μL/well of various DNA antigens (2 μg/mL) diluted in Saline-Sodium Citrate (SSC) Buffer. Control wells had SSC alone. Coated plates were incubated overnight at 4 °C. The next day, plates were washed with PBS, followed by blocking for 2 hours at room temperature with blocking buffer (2% bovine serum albumin (BSA), 0.05% Tween-20 in PBS). After blocking, the plates were washed with PBS and incubated for 1 hour at room temperature with horseradish peroxidase (HRP)-conjugated secondary reagent: anti-mouse IgG (γ chain specific [EMD Millipore]) at 1:1000; anti-sheep IgG (H + L chain specific [EMD Millipore]) at 1:1000; or anti-human IgG (γ chain specific [Sigma-Aldrich]) at 1:1500. The reaction proceeded for 1 hour at room temperature. The secondary antibodies were diluted with PBS ELISA dilution buffer (P-EDB; 0.1% BSA, 0.05% Tween 20 in PBS, pH 7.4). The plates were washed, then incubated with TMB substrate for 30 minutes at room temperature. Sulfuric acid was then added to terminate the color development. The absorbance was measured at 450 nm using a UVmax multi-plate spectrophotometer (Molecular Devices).

### Infection of mice

6-8 week-old 129X1/SvJ mice, including both males and females, were inoculated intragastrically with 20 mg of streptomycin (0.1ml of a 200 mg/mL solution in PBS) 24 hours before bacterial inoculation to induce gastrointestinal pathology [82]. Bacteria were grown shaking in LB broth at 37°C overnight. Mice were inoculated intragastrically with either 0.1 mL of sterile LB broth (mock infection) or 10^7^−10^8^ CFU *S.* Typhimurium IR715 or the *invAspiB* mutant.

For LD experiments, 6-8 week old male 129X1/SvJ mice were fed a high cholesterol (1% cholesterol, 0.5% cholic acid, Envigo, TD 140673; lithogenic) diet for 6 weeks. After this time, mice were taken off the high-cholesterol diet and put on a normal diet for 10 days. Mice were infected i.p. with 2 × 10^3^ STm strain 14028, without streptomycin pre-treatment. For the Z-DNA staining in cecal tissue, 6-8 weeks old male and female 129X1/SvJ mice were fed a lithogenic diet and then orally infected with 1 × 10^^3^ STm 14028 for 3 weeks. Cecal tissue was collected, fixed in 10% formalin for 48 hours, then paraffin-embedded, and tissue sections were stained as previously described[74]. Cecal tissue sections were stained with primary antibodies rabbit *Salmonella* O antiserum (1:500, Difco, 226591) and mouse anti-Z-DNA (1:200, Z22, Absolute Antibodies) or EBBA Biolight 680 bacterial amyloid dye (1:200), and secondary antibodies AlexaFluor488-conjugated goat anti-rabbit IgG (1:250, Life Technologies, A11008) and Rhodamine Red X-conjugated AffiniPure donkey anti-mouse IgG (H+L) (1:100, Jackson Immuno Research, 715-295-150). Sections were imaged at 63X magnification on the Leica SP5 confocal microscope and images were processed on ImageJ2.

### Z-HUNT algorithm

To evaluate the thermodynamic propensity of genomic regions to adopt the Z-conformation, we employed the Z-Hunt[rs] (ZH) program (v0.0.4, https://github.com/biomancy/zhuntrs), a Rust-based reimplementation of the original Z-Hunt algorithm[83] (v3), together with Z-DNABERT, a deep-learning model trained to identify Z-DNA-prone sequence contexts[84]. All available annotated type genomes for each species were obtained from NCBI GenBank, and “ZH-scores” were predicted across each chromosome using fixed-size windows (12–15 bp; corresponding to 6–7 dinucleotides). For the scatter plot visualization of selected bacterial and mitochondrial genomes, GC content was calculated as the percentage of guanine (G) and cytosine (C) bases in the total genome. Z-DNA site density was quantified as the number of Z-Hunt predicted sites per kilobase (sites/kb). Spearman rank correlation was calculated to assess the relationship between GC content and Z-DNA site density across all genome assemblies.

For Circos plot visualization of selected genomes, Z-DNABERT was run in conjunction with ZHUNT using the publicly available HG-Kouzine model to filter for highly Z-prone sequences. The genome was scanned in small overlapping 6-base segments, and the model’s predicted Z-DNA scores were compiled across the chromosome. Regions with consistently high scores (≥0.9) were marked and saved, and only stretches at least 12 bp long were retained as Z-prone candidates.

### STm lysis and nucleic acid staining

A 5 mL overnight STm culture in LB supplemented with 50 μg/mL nalidixic acid was pelleted by centrifugation at 4000 x g for 5 minutes, resuspended in 1 mL 4% formaldehyde in PBS, and incubated at RT for 30 minutes. The pellet was then washed three times in PBS. After the final centrifugation, the supernatant was removed, and the pellet was resuspended in 70% ethanol and incubated for 1 hour at room temperature with shaking. To immobilize cells, an 8-well slide was coated with poly-L-lysine for 1 hour and washed three times with PBS prior to addition of cells. Cells were mixed with 1:1 volume water, centrifuged at 600 x g for 5 minutes, resuspended in PBS and mounted to pre-coated slide wells. Cells were incubated with 1 mg/mL lysozyme in Tris-EDTA-Glycine buffer for 30 minutes at room temperature. Staining and imaging of the nucleic acids were performed as described.

### STm genomic dna antibody specificity

STm genomic DNA was extracted following the instructions of the E.Z.N.A Bacterial DNA Extraction Kit (Thomas Scientific, C755C96). Antibody specificity was determined via ELISA, as described above.

### Statistical analysis

Data were analyzed using Prism software (GraphPad). Two-way ANOVA with post-hoc Tukey multiple comparisons tests or two-tailed Student’s t test were used as appropriate. The p-values <0.05 were considered significant. *p < 0.05, **p < 0.01, ***p < 0.001 were marked in the figures.

## Ethics statement

Experiments were performed under protocols approved by AAALAC-accredited Temple University Lewis Katz School of Medicine and Nationwide Children’s Hospital, Institutional Animal Care and Use Committee (IACUC), on file with the NIH Office for the Protection of Research Risks, in accordance with USDA and PHS Policy on Humane Care and Use of Laboratory Animals.

## ACKNOWLEDGEMENTS

We would like to thank Drs. Steven Goodman (Nationwide Children’s Hospital), Marc Monestier (Tempe University Lewis Katz School of Medicine), Alex Fedorov (University of Oxford) and Stefania Gallucci (University of Massachusetts) for their assistance with this study.

## REFERENCES

1. Justiz Vaillant AA, Goyal A, Varacallo MA. Systemic Lupus Erythematosus. In: StatPearls [Internet]. Treasure Island (FL): StatPearls Publishing; 2025 [cited 2026 Jan 13]. Available from: http://www.ncbi.nlm.nih.gov/books/NBK535405/ PubMed PMID: 30571026.

2. Hahn BH. Antibodies to DNA. N Engl J Med. 1998 May 7;338(19):1359–68. doi:10.1056/NEJM199805073381906

3. Pisetsky DS. Anti-DNA antibodies--quintessential biomarkers of SLE. Nat Rev Rheumatol. 2016 Feb;12(2):102–10. doi:10.1038/nrrheum.2015.151 PubMed PMID: 26581343.

4. Jang YJ, Stollar BD. Anti-DNA antibodies: aspects of structure and pathogenicity. Cell Mol Life Sci CMLS. 2003 Feb;60(2):309–20. doi:10.1007/s000180300026 PubMed PMID: 12678496; PubMed Central PMCID: PMC11138676.

5. Rich A, Nordheim A, Wang AH. The chemistry and biology of left-handed Z-DNA. Annu Rev Biochem. 1984;53:791–846. doi:10.1146/annurev.bi.53.070184.004043 PubMed PMID: 6383204.

6. Herbert A. Z-DNA and Z-RNA in human disease. Commun Biol. 2019 Jan 7;2(1):7. doi:10.1038/s42003-018-0237-x

7. Azorin F, Nordheim A, Rich A. Formation of Z-DNA in negatively supercoiled plasmids is sensitive to small changes in salt concentration within the physiological range. EMBO J. 1983 May 1;2(5):649–55. doi:10.1002/j.1460-2075.1983.tb01479.x

8. Herbert A. ALU non-B-DNA conformations, flipons, binary codes and evolution. R Soc Open Sci. 2020 Jun 3;7(6):200222. doi:10.1098/rsos.200222

9. Ho PS. Thermogenomics: thermodynamic-based approaches to genomic analyses of DNA structure. Methods San Diego Calif. 2009 Mar;47(3):159–67. doi:10.1016/j.ymeth.2008.09.007 PubMed PMID: 18848994.

10. Moller A, Nordheim A, Kozlowski SA, Patel D, Rich A. Bromination stabilizes poly(dG-dC) in the Z-DNA form under low-salt conditions. Biochemistry. 1984 Jan 3;23(1):54–62. doi:10.1021/bi00296a009

11. Hanau LH, Santella RM, Grunberger D, Erlanger BF. An immunochemical examination of acetylaminofluorene-modified poly(dG-dC) X poly(dG-dC) in the Z-conformation. J Biol Chem. 1984 Jan 10;259(1):173–8. doi:10.1016/S0021-9258(17)43637-4

12. Lafer EM, Möller A, Nordheim A, Stollar BD, Rich A. Antibodies specific for left-handed Z-DNA. Proc Natl Acad Sci. 1981 Jun;78(6):3546–50. doi:10.1073/pnas.78.6.3546

13. Lafer EM, Moller A, Valle RPC, Nordheim A, Rich A, Stollar BD. Antibody recognition of Z-DNA. Unkn J. 1982;47(1):155–62. doi:10.1101/sqb.1983.047.01.020

14. Möller A, Gabriels JE, Lafer EM, Nordheim A, Rich A, Stollar BD. Monoclonal antibodies recognize different parts of Z-DNA. J Biol Chem. 1982 Oct 25;257(20):12081–5. doi:10.1016/S0021-9258(18)33681-0

15. Edgington SM, Stollar BD. Immunogenicity of Z-DNA depends on the size of polynucleotide presented in complexes with methylated BSA. Mol Immunol. 1992 May 1;29(5):609–17. doi:10.1016/0161-5890(92)90197-6

16. Stollar BD. Why the difference between B-DNA and Z-DNA? Lupus. 1997;6(3):327–8. doi:10.1177/096120339700600327 PubMed PMID: 9296781.

17. Tursi SA, Tükel Ç. Curli-Containing Enteric Biofilms Inside and Out: Matrix Composition, Immune Recognition, and Disease Implications. Microbiol Mol Biol Rev. 2018 Oct 10;82(4):10.1128/mmbr.00028-18. doi:10.1128/mmbr.00028-18

18. Miller AL, Pasternak JA, Medeiros NJ, Nicastro LK, Tursi SA, Hansen EG, et al. In vivo synthesis of bacterial amyloid curli contributes to joint inflammation during S. Typhimurium infection. PLOS Pathog. 2020 Jul 9;16(7):e1008591. doi:10.1371/journal.ppat.1008591

19. Crawford RW, Gibson DL, Kay WW, Gunn JS. Identification of a bile-induced exopolysaccharide required for Salmonella biofilm formation on gallstone surfaces. Infect Immun. 2008 Nov;76(11):5341–9. doi:10.1128/IAI.00786-08 PubMed PMID: 18794278; PubMed Central PMCID: PMC2573354.

20. Cruz-Cruz A, Schreeg ME, Gunn JS. A temporary cholesterol-rich diet and bacterial extracellular matrix factors favor Salmonella spp. biofilm formation in the cecum. mBio. 2024 Dec 5;16(1):e03242–24. doi:10.1128/mbio.03242-24

21. Albicoro FJ, Bessho S, Grando K, Olubajo S, Tam V, Tükel Ç. Lactate promotes the biofilm-to-invasive-planktonic transition in Salmonella enterica serovar Typhimurium via the de novo purine pathway. Infect Immun. 92(10):e00266–24. doi:10.1128/iai.00266-24 PubMed PMID: 39133016; PubMed Central PMCID: PMC11475809.

22. Miller AL, Nicastro LK, Bessho S, Grando K, White AP, Zhang Y, et al. Nitrate Is an Environmental Cue in the Gut for Salmonella enterica Serovar Typhimurium Biofilm Dispersal through Curli Repression and Flagellum Activation via Cyclic-di-GMP Signaling. mBio. 2021 Feb 22;13(1):e0288621. doi:10.1128/mbio.02886-21 PubMed PMID: 35130730; PubMed Central PMCID: PMC8822344.

23. Tursi SA, Lee EY, Medeiros NJ, Lee MH, Nicastro LK, Buttaro B, et al. Bacterial amyloid curli acts as a carrier for DNA to elicit an autoimmune response via TLR2 and TLR9. PLoS Pathog. 2017 Apr 14;13(4):e1006315. doi:10.1371/journal.ppat.1006315 PubMed PMID: 28410407; PubMed Central PMCID: PMC5406031.

24. Gallo PM, Rapsinski GJ, Wilson RP, Oppong GO, Sriram U, Goulian M, et al. Amyloid-DNA Composites of Bacterial Biofilms Stimulate Autoimmunity. Immunity. 2015 Jun 16;42(6):1171–84. doi:10.1016/j.immuni.2015.06.002

25. Issac JM, Mohamed YA, Bashir GH, Al-Sbiei A, Conca W, Khan TA, et al. Induction of Hypergammaglobulinemia and Autoantibodies by Salmonella Infection in MyD88-Deficient Mice. Front Immunol. 2018 Jun 20;9. doi:10.3389/fimmu.2018.01384

26. Gallo PM, Rapsinski GJ, Wilson RP, Oppong GO, Sriram U, Goulian M, et al. Amyloid-DNA composites of bacterial biofilms stimulate autoimmunity. Immunity. 2015 Jun 16;42(6):1171–84. doi:10.1016/j.immuni.2015.06.002 PubMed PMID: 26084027; PubMed Central PMCID: PMC4500125.

27. Nicastro LK, de Anda J, Jain N, Grando KCM, Miller AL, Bessho S, et al. Assembly of ordered DNA-curli fibril complexes during Salmonella biofilm formation correlates with strengths of the type I interferon and autoimmune responses. PLoS Pathog. 2022 Aug 16;18(8):e1010742. doi:10.1371/journal.ppat.1010742 PubMed PMID: 35972973; PubMed Central PMCID: PMC9380926.

28. Bessho S, Grando KCM, Kyrylchuk K, Miller A, Klein-Szanto AJ, Zhu W, et al. Systemic exposure to bacterial amyloid curli alters the gut mucosal immune response and the microbiome, exacerbating Salmonella-induced arthritis. Gut Microbes. 15(1):2221813. doi:10.1080/19490976.2023.2221813 PubMed PMID: 37317012; PubMed Central PMCID: PMC10269392.

29. Grando K, Bessho S, Harrell K, Kyrylchuk K, Pantoja AM, Olubajo S, et al. Bacterial amyloid curli activates the host unfolded protein response via IRE1α in the presence of HLA-B27. Gut Microbes. 2024;16(1):2392877. doi:10.1080/19490976.2024.2392877 PubMed PMID: 39189642; PubMed Central PMCID: PMC11352795.

30. González JF, Tucker L, Fitch J, Wetzel A, White P, Gunn JS. Human Bile-Mediated Regulation of Salmonella Curli Fimbriae. J Bacteriol. 2019 Aug 22;201(18):10.1128/jb.00055-19. doi:10.1128/jb.00055-19

31. Crawford RW, Reeve KE, Gunn JS. Flagellated but Not Hyperfimbriated Salmonella enterica Serovar Typhimurium Attaches to and Forms Biofilms on Cholesterol-Coated Surfaces. J Bacteriol. 2010 Jun;192(12):2981–90. doi:10.1128/JB.01620-09 PubMed PMID: 20118264; PubMed Central PMCID: PMC2901699.

32. Buzzo JR, Devaraj A, Gloag ES, Jurcisek JA, Robledo-Avila F, Kesler T, et al. Z-form extracellular DNA is a structural component of the bacterial biofilm matrix. Cell. 2021 Nov;184(23):5740–5758.e17. doi:10.1016/j.cell.2021.10.010

33. Ramesh N, Brahmachari SK. Structural alteration from non-B to B-form could reflect DNase I hypersensitivity. J Biomol Struct Dyn. 1989 Apr;6(5):899–906. doi:10.1080/07391102.1989.10506521 PubMed PMID: 2590508.

34. Bhanjadeo MM, Nayak AK, Subudhi U. Cerium chloride stimulated controlled conversion of B-to-Z DNA in self-assembled nanostructures. Biochem Biophys Res Commun. 2017 Jan 22;482(4):916–21. doi:10.1016/j.bbrc.2016.11.133 PubMed PMID: 27890616.

35. Kwakye-Berko F, Meshnick S. Sequence preference of chloroquine binding to DNA and prevention of Z-DNA formation. Mol Biochem Parasitol. 1990 Mar;39(2):275–8. doi:10.1016/0166-6851(90)90066-u PubMed PMID: 2320060.

36. Amar Y, Lagkouvardos I, Silva RL, Ishola OA, Foesel BU, Kublik S, et al. Pre-digest of unprotected DNA by Benzonase improves the representation of living skin bacteria and efficiently depletes host DNA. Microbiome. 2021 May 26;9:123. doi:10.1186/s40168-021-01067-0 PubMed PMID: 34039428; PubMed Central PMCID: PMC8157445.

37. Nestle M, Roberts WK. An extracellular nuclease from Serratia marcescens. II. Specificity of the enzyme. J Biol Chem. 1969 Oct 10;244(19):5219–25. PubMed PMID: 4310088.

38. Hung C, Zhou Y, Pinkner JS, Dodson KW, Crowley JR, Heuser J, et al. Escherichia coli biofilms have an organized and complex extracellular matrix structure. mBio. 2013 Sep 10;4(5):e00645–00613. doi:10.1128/mBio.00645-13 PubMed PMID: 24023384; PubMed Central PMCID: PMC3774191.

39. Spencer DM, Reyna AG, Pisetsky DS. The Binding of Monoclonal and Polyclonal Anti-Z-DNA Antibodies to DNA of Various Species Origin. Int J Mol Sci. 2021 Jan;22(16):8931. doi:10.3390/ijms22168931

40. Pisetsky DS, Gedye MJ, David LA, Spencer DM. The Binding Properties of Antibodies to Z-DNA in the Sera of Normal Healthy Subjects. Int J Mol Sci. 2024 Feb 22;25(5):2556. doi:10.3390/ijms25052556 PubMed PMID: 38473808; PubMed Central PMCID: PMC10931986.

41. Schroth GP, Chou PJ, Ho PS. Mapping Z-DNA in the human genome. Computer-aided mapping reveals a nonrandom distribution of potential Z-DNA-forming sequences in human genes. J Biol Chem. 1992 Jun 15;267(17):11846–55. PubMed PMID: 1601856.

42. Jarvik T, Smillie C, Groisman EA, Ochman H. Short-Term Signatures of Evolutionary Change in the Salmonella enterica Serovar Typhimurium 14028 Genome. J Bacteriol. 2010 Jan;192(2):560–7. doi:10.1128/JB.01233-09 PubMed PMID: 19897643; PubMed Central PMCID: PMC2805332.

43. Papanikolaou N, Trachana K, Theodosiou T, Promponas VJ, Iliopoulos I. Gene socialization: gene order, GC content and gene silencing in Salmonella. BMC Genomics. 2009 Dec 11;10:597. doi:10.1186/1471-2164-10-597 PubMed PMID: 20003346; PubMed Central PMCID: PMC2801525.

44. Piovesan A, Pelleri MC, Antonaros F, Strippoli P, Caracausi M, Vitale L. On the length, weight and GC content of the human genome. BMC Res Notes. 2019 Feb 27;12(1):106. doi:10.1186/s13104-019-4137-z PubMed PMID: 30813969; PubMed Central PMCID: PMC6391780.

45. Azzouz D, Omarbekova A, Heguy A, Schwudke D, Gisch N, Rovin BH, et al. Lupus nephritis is linked to disease-activity associated expansions and immunity to a gut commensal. Ann Rheum Dis. 2019 Jul;78(7):947–56. doi:10.1136/annrheumdis-2018-214856 PubMed PMID: 30782585; PubMed Central PMCID: PMC6585303.

46. Pisetsky DS, Garza Reyna A, Belina ME, Spencer DM. The Interaction of Anti-DNA Antibodies with DNA: Evidence for Unconventional Binding Mechanisms. Int J Mol Sci. 2022 May 7;23(9):5227. doi:10.3390/ijms23095227 PubMed PMID: 35563617; PubMed Central PMCID: PMC9105193.

47. Carlé C, Fortenfant F, Bost C, Belliere J, Faguer S, Chauveau D, et al. The added value of coupling anti-dsDNA and anti-chromatin antibodies in follow-up monitoring of systemic lupus erythematosus patients. J Transl Autoimmun. 2025 Jan 22;10:100274. doi:10.1016/j.jtauto.2025.100274 PubMed PMID: 39917317; PubMed Central PMCID: PMC11799958.

48. Schroeder K, Wellmann U, Winkler TH, Herrmann M. Evolution of anti-DNA autoantibodies by somatic hypermutation: evidence for postmutational B cell tolerance. Ann Rheum Dis. 2012 Feb 1;71:A32. doi:10.1136/annrheumdis-2011-201234.1

49. Mevorach D, Zhou JL, Song X, Elkon KB. Systemic Exposure to Irradiated Apoptotic Cells Induces Autoantibody Production. J Exp Med. 1998 Jul 20;188(2):387–92. doi:10.1084/jem.188.2.387 PubMed PMID: 9670050; PubMed Central PMCID: PMC2212450.

50. Mahajan A, Herrmann M, Muñoz LE. Clearance Deficiency and Cell Death Pathways: A Model for the Pathogenesis of SLE. Front Immunol. 2016 Feb 8;7. doi:10.3389/fimmu.2016.00035

51. Azzouz DF, Chen Z, Izmirly PM, Chen LA, Li Z, Zhang C, et al. Longitudinal gut microbiome analyses and blooms of pathogenic strains during lupus disease flares. Ann Rheum Dis. 2023 Oct;82(10):1315–27. doi:10.1136/ard-2023-223929 PubMed PMID: 37365013; PubMed Central PMCID: PMC10511964.

52. He J, Chan T, Hong X, Zheng F, Zhu C, Yin L, et al. Microbiome and Metabolome Analyses Reveal the Disruption of Lipid Metabolism in Systemic Lupus Erythematosus. Front Immunol. 2020 Jul 31;11:1703. doi:10.3389/fimmu.2020.01703 PubMed PMID: 32849599; PubMed Central PMCID: PMC7411142.

53. Silverman GJ, Deng J, Azzouz DF. Sex-dependent Lupus Blautia (Ruminococcus) gnavus strain induction of zonulin-mediated intestinal permeability and autoimmunity. Front Immunol. 2022 Aug 11;13:897971. doi:10.3389/fimmu.2022.897971 PubMed PMID: 36032126; PubMed Central PMCID: PMC9405438.

54. Rogers JV, Hall VL, McOsker CC. Crumbling the Castle: Targeting DNABII Proteins for Collapsing Bacterial Biofilms as a Therapeutic Approach to Treat Disease and Combat Antimicrobial Resistance. Antibiot Basel Switz. 2022 Jan 14;11(1):104. doi:10.3390/antibiotics11010104 PubMed PMID: 35052981; PubMed Central PMCID: PMC8773079.

55. Devaraj A, Buzzo JR, Mashburn-Warren L, Gloag ES, Novotny LA, Stoodley P, et al. The extracellular DNA lattice of bacterial biofilms is structurally related to Holliday junction recombination intermediates. Proc Natl Acad Sci U S A. 2019 Dec 10;116(50):25068–77. doi:10.1073/pnas.1909017116 PubMed PMID: 31767757; PubMed Central PMCID: PMC6911203.

56. Devaraj A, Justice SS, Bakaletz LO, Goodman SD. DNABII proteins play a central role in UPEC biofilm structure. Mol Microbiol. 2015 Jun;96(6):1119–35. doi:10.1111/mmi.12994 PubMed PMID: 25757804; PubMed Central PMCID: PMC4464964.

57. Goodman SD, Obergfell KP, Jurcisek JA, Novotny LA, Downey JS, Ayala EA, et al. Biofilms can be dispersed by focusing the immune system on a common family of bacterial nucleoid-associated proteins. Mucosal Immunol. 2011 Nov;4(6):625–37. doi:10.1038/mi.2011.27 PubMed PMID: 21716265.

58. Tursi SA, Lee EY, Medeiros NJ, Lee MH, Nicastro LK, Buttaro B, et al. Bacterial amyloid curli acts as a carrier for DNA to elicit an autoimmune response via TLR2 and TLR9. PLoS Pathog. 2017 Apr 14;13(4):e1006315. doi:10.1371/journal.ppat.1006315 PubMed PMID: 28410407; PubMed Central PMCID: PMC5406031.

59. Tükel Ç, Wilson RP, Nishimori JH, Pezeshki M, Chromy BA, Bäumler AJ. Responses to Amyloids of Microbial and Host Origin Are Mediated through Toll-like Receptor 2. Cell Host Microbe. 2009 Jul 23;6(1):45–53. doi:10.1016/j.chom.2009.05.020 PubMed PMID: 19616765.

60. Tükel C, Nishimori JH, Wilson RP, Winter MG, Keestra AM, van Putten JPM, et al. Toll-like receptors 1 and 2 cooperatively mediate immune responses to curli, a common amyloid from enterobacterial biofilms. Cell Microbiol. 2010 Oct;12(10):1495–505. doi:10.1111/j.1462-5822.2010.01485.x PubMed PMID: 20497180; PubMed Central PMCID: PMC3869100.

61. Rapsinski GJ, Wynosky-Dolfi MA, Oppong GO, Tursi SA, Wilson RP, Brodsky IE, et al. Toll-like receptor 2 and NLRP3 cooperate to recognize a functional bacterial amyloid, curli. Infect Immun. 2015 Feb;83(2):693–701. doi:10.1128/IAI.02370-14 PubMed PMID: 25422268; PubMed Central PMCID: PMC4294241.

62. Tükel C, Raffatellu M, Humphries AD, Wilson RP, Andrews-Polymenis HL, Gull T, et al. CsgA is a pathogen-associated molecular pattern of Salmonella enterica serotype Typhimurium that is recognized by Toll-like receptor 2. Mol Microbiol. 2005 Oct;58(1):289–304. doi:10.1111/j.1365-2958.2005.04825.x PubMed PMID: 16164566.

63. Forero-Peña DA, Carrión-Nessi FS, Lopez-Perez M, Sandoval-de Mora M, Amaya ID, Gamardo ÁF, et al. Seroprevalence of viral and bacterial pathogens among malaria patients in an endemic area of southern Venezuela. Infect Dis Poverty. 2023 Apr 10;12:33. doi:10.1186/s40249-023-01089-w PubMed PMID: 37038195; PubMed Central PMCID: PMC10084699.

64. Fine RL, Manfredo Vieira S, Gilmore MS, Kriegel MA. Mechanisms and consequences of gut commensal translocation in chronic diseases. Gut Microbes. 2019 Jul 15;11(2):217–30. doi:10.1080/19490976.2019.1629236 PubMed PMID: 31306081; PubMed Central PMCID: PMC7053960.

65. Manfredo Vieira S, Hiltensperger M, Kumar V, Zegarra-Ruiz D, Dehner C, Khan N, et al. Translocation of a gut pathobiont drives autoimmunity in mice and humans. Science. 2018 Mar 9;359(6380):1156–61. doi:10.1126/science.aar7201 PubMed PMID: 29590047; PubMed Central PMCID: PMC5959731.

66. Hill Gaston JS, Lillicrap MS. Arthritis associated with enteric infection. Best Pract Res Clin Rheumatol. 2003 Apr;17(2):219–39. doi:10.1016/s1521-6942(02)00104-3 PubMed PMID: 12787523.

67. Durand DV, Lecomte C, Cathébras P, Rousset H, Godeau P, Disease the SRG on W. Whipple Disease: Clinical Review of 52 Cases. Medicine (Baltimore). 1997 May;76(3):170.

68. Kocar ZC kaner S Pay, and M Turan, IH. Clostridium Difficile Infection in Patients with Reactive Arthritis of Undetermined Etiology. Scand J Rheumatol. 1998 Jan 1;27(5):357–62. doi:10.1080/03009749850154384 PubMed PMID: 9808399.

69. Noer HR. An “Experimental” Epidemic of Reiter’s Syndrome. JAMA. 1966 Nov 14;198(7):693–8. doi:10.1001/jama.1966.03110200049016

70. Hannu T, Mattila L, Rautelin H, Pelkonen P, Lahdenne P, Siitonen A, et al. Campylobacter-triggered reactive arthritis: a population-based study. Rheumatology. 2002 Mar 1;41(3):312–8. doi:10.1093/rheumatology/41.3.312

71. Dworkin MS, Shoemaker PC, Goldoft MJ, Kobayashi JM. Reactive Arthritis and Reiter’s Syndrome Following an Outbreak of Gastroenteritis Caused by Salmonella enteritidis. Clin Infect Dis. 2001 Oct 1;33(7):1010–4. doi:10.1086/322644

72. Fendler C, Laitko S, Sörensen H, Gripenberg-Lerche C, Groh A, Uksila J, et al. Frequency of triggering bacteria in patients with reactive arthritis and undifferentiated oligoarthritis and the relative importance of the tests used for diagnosis. Ann Rheum Dis. 2001 Apr 1;60(4):337–43. doi:10.1136/ard.60.4.337

73. Laitio P, Virtala M, Salmi M, Pelliniemi LJ, Yu DT, Granfors K. HLA-B27 modulates intracellular survival of Salmonella enteritidis in human monocytic cells. Eur J Immunol. 1997 Jun;27(6):1331–8. doi:10.1002/eji.1830270606 PubMed PMID: 9209481.

74. Grando K, Bessho S, Harrell K, Kyrylchuk K, Pantoja AM, Olubajo S, et al. Bacterial amyloid curli activates the host unfolded protein response via IRE1α in the presence of HLA-B27. Gut Microbes. 2024;16(1):2392877. doi:10.1080/19490976.2024.2392877 PubMed PMID: 39189642; PubMed Central PMCID: PMC11352795.

75. Mattila L, Leirisalo-Repo M, Pelkonen P, Koskimies S, Granfors K, Siitonen A. Reactive arthritis following an outbreak of Salmonella Bovismorbificans infection. J Infect. 1998 May;36(3):289–95. doi:10.1016/s0163-4453(98)94243-8 PubMed PMID: 9661939.

76. Ekman P, Kirveskari J, Granfors K. Modification of disease outcome in Salmonella-infected patients by HLA-B27. Arthritis Rheum. 2000 Jul;43(7):1527–34. doi:10.1002/1529-0131(200007)43:7%3C1527::AID-ANR17%3E3.0.CO;2-G PubMed PMID: 10902756.

77. Stojiljkovic I, Bäumler AJ, Heffron F. Ethanolamine utilization in Salmonella typhimurium: nucleotide sequence, protein expression, and mutational analysis of the cchA cchB eutE eutJ eutG eutH gene cluster. J Bacteriol. 1995 Mar;177(5):1357–66. doi:10.1128/jb.177.5.1357-1366.1995 PubMed PMID: 7868611; PubMed Central PMCID: PMC176743.

78. Raffatellu M, Wilson RP, Chessa D, Andrews-Polymenis H, Tran QT, Lawhon S, et al. SipA, SopA, SopB, SopD, and SopE2 contribute to Salmonella enterica serotype typhimurium invasion of epithelial cells. Infect Immun. 2005 Jan;73(1):146–54. doi:10.1128/IAI.73.1.146-154.2005 PubMed PMID: 15618149; PubMed Central PMCID: PMC538951.

79. Nishimori JH, Newman TN, Oppong GO, Rapsinski GJ, Yen JH, Biesecker SG, et al. Microbial Amyloids Induce Interleukin 17A (IL-17A) and IL-22 Responses via Toll-Like Receptor 2 Activation in the Intestinal Mucosa. Infect Immun. 2012 Nov 12;80(12):4398–408. doi:10.1128/iai.00911-12

80. Nicastro LK, Tursi SA, Le LS, Miller AL, Efimov A, Buttaro B, et al. Cytotoxic Curli Intermediates Form during Salmonella Biofilm Development. J Bacteriol. 2019 Sep 15;201(18):e00095–19. doi:10.1128/JB.00095-19 PubMed PMID: 31182496; PubMed Central PMCID: PMC6707925.

81. Spencer DM, Svenungsson E, Gunnarsson I, Caricchio R, Pisetsky DS. The expression of antibodies to Z-DNA in the blood of patients with systemic lupus erythematosus: Relationship to autoantibodies to B-DNA. Clin Immunol Orlando Fla. 2023 Oct;255:109763. doi:10.1016/j.clim.2023.109763 PubMed PMID: 37673226.

82. Barthel M, Hapfelmeier S, Quintanilla-Martínez L, Kremer M, Rohde M, Hogardt M, et al. Pretreatment of Mice with Streptomycin Provides a Salmonella enterica Serovar Typhimurium Colitis Model That Allows Analysis of Both Pathogen and Host. Infect Immun. 2003 May;71(5):2839–58. doi:10.1128/IAI.71.5.2839-2858.2003 PubMed PMID: 12704158; PubMed Central PMCID: PMC153285.

83. Ho PS, Ellison MJ, Quigley GJ, Rich A. A computer aided thermodynamic approach for predicting the formation of Z-DNA in naturally occurring sequences. EMBO J. 1986 Oct;5(10):2737–44. doi:10.1002/j.1460-2075.1986.tb04558.x PubMed PMID: 3780676; PubMed Central PMCID: PMC1167176.

84. Umerenkov D, Herbert A, Konovalov D, Danilova A, Beknazarov N, Kokh V, et al. Z-flipon variants reveal the many roles of Z-DNA and Z-RNA in health and disease. Life Sci Alliance. 2023 Jul;6(7):e202301962. doi:10.26508/lsa.202301962 PubMed PMID: 37164635; PubMed Central PMCID: PMC10172764.

